# Time as a supervisor: temporal regularity and auditory object learning

**DOI:** 10.1101/2022.11.10.515986

**Authors:** Ronald W. Di Tullio, Chetan Parthiban, Eugenio Piasini, Pratik Chaudhari, Vijay Balasubramanian, Yale E. Cohen

## Abstract

Sensory systems appear to learn to transform incoming sensory information into perceptual representations, or “objects”, that can inform and guide behavior with minimal explicit supervision. Here, we propose that the auditory system can achieve this goal by using time as a supervisor, i.e., by learning features of a stimulus that are temporally regular. We will show that this procedure generates a feature space sufficient to support fundamental computations of auditory perception. In detail, we consider the problem of discriminating between instances of a prototypical class of natural auditory objects, i.e., rhesus macaque vocalizations. We test discrimination in two ethologically relevant tasks: discrimination in a cluttered acoustic background and generalization to discriminate between novel exemplars. We show that an algorithm that learns these temporally regular features affords better or equivalent discrimination and generalization than conventional feature-selection algorithms, i.e., principal component analysis and independent component analysis. Our findings suggest that the slow temporal features of auditory stimuli may be sufficient for parsing auditory scenes and that the auditory brain could utilize these slowly changing temporal features.

---

Hearing animals parse their auditory world into distinct perceptual representations (i.e., auditory objects.) that inform and guide behavior. To do so, the auditory system learns to group stimuli with similar spectrotemporal properties into one perceptual object, while simultaneously segregating stimuli with different properties into a different object or objects [1–7]. This learning task is complicated because natural objects can have the same identity despite large variations in physical properties, e.g., their location [8–12]; they are often encountered in complex, cluttered environments, e.g., noisy rooms filled with speakers [13–18]; and it is often necessary to generalize to novel objects, e.g., the speech of new acquaintances whose voices have different timbres, accents, and pitches [3, 6, 19]. Despite these challenges, the auditory system appears to learn this stimulus-to-object transformation with minimal explicit supervision.

Intuitively, objects are entities composed of correlated components that change together, so that the whole retains its coherence and identity, even in the presence of other variations [20]. If natural objects are defined in this way, it should be possible to learn and discriminate between them based on temporal cues including: 1) coincidence between the components, 2) continuity in the components, and 3) continuity in the correlations between components. That is, the temporal regularities of an object should define its identity.

This notion that an object is defined by its temporal properties is consistent with our current interpretation of the Gestalt principles underlying auditory scene analysis [1, 21]. One of these principles suggests how temporal sequences of stimuli are assigned to common sources or objects. For example, although we hear the sound that occurs when someone steps on the ground, we bind the temporal series of such sounds into “footsteps”. Likewise, we can segregate footsteps of someone running in sneakers from someone strolling in wooden clogs, based not only differences in their frequency structure but also on temporal properties like the relative timing between each step. Such Gestalt principles [1, 21] provide important intuitions for what defines an auditory object and how the auditory system may discriminate between objects. However, they do not provide a unified computational principle that explains how such temporal information could actually be learned and utilized for discrimination by the auditory system. Here, we seek to provide such a computational understanding.

We use rhesus macaque vocalizations as prototypical natural auditory objects [22] because their acoustic structure is similar to that seen in human speech, bird song, and other animal vocalizations [23, 24]. We replicate previous results showing these vocalizations are defined by having temporally continuous components. We then expand on these results by showing this structure extends to the continuity of correlations between components and that this structure is significantly less present or virtually absent for other classes of stimuli.

Next, we use Slow Feature Analysis (SFA), a temporal learning algorithm originally used to study vision [25], to infer auditory features from four different acoustic classes of macaque vocalizations as well as control stimuli. For our purposes, SFA serves as a convenient method for extracting linear and nonlinear features [26] that summarizes the most temporally continuous components and correlations between components of auditory stimuli, respectively.

Finally, we train linear support vector machine (SVM) classifiers to separate vocalizations in low-dimensional subspaces of the learned SFA feature space. Our SFA-based classifiers outperform Principal Component Analysis (PCA) and Independent Component Analysis (ICA)-based classifiers, which have no explicit notion temporal regularity but instead respectively assume that the most variable features or the features that generate the most statistically independent outputs are informative about object identity. We confirm consistency with our hypothesis: our classifiers perform at chance on white noise, and at intermediate levels on applause, an auditory texture [11, 27] generated by the superposition of claps. These results also hold after addition of clutter to the training and test sets, showing that SFA-based classifiers can solve the “cocktail party problem”. Finally, our classifiers generalize to novel exemplars of all four vocalization classes.

Our results suggest that time can supervise the learning of natural auditory objects. That is, auditory features capturing the most temporally continuous components and correlations of auditory stimuli may suffice for parsing auditory scenes, implying that the brain could be tuned to these slowly changing temporal features.

## I. RESULTS

### A. Temporal regularity in natural auditory stimuli

We propose that auditory objects are fundamentally defined by temporal regularity, specifically through the continuity of components of the acoustic signal and of correlations between these components. (Fig. 1A). Thus, we first tested whether these continuities and correlations can be seen directly in auditory stimuli. To do so, we had to select between two broad categories of natural auditory stimuli: acoustic events, such as vocalizations, and textures, such as rain [3, 6, 11, 27]. We focused on acoustic events, exemplified by rhesus macaque vocalizations, because of their ethological relevance [22], prototypical structure [23, 24], and links to previous work on neural correlates of auditory perception [4, 6, 24]. By contrast, textures appear to be defined by longer-time scale/time averaged statistics [11, 27], and thus it is possible the brain processes them differently than acoustic events.

**Figure 1:**
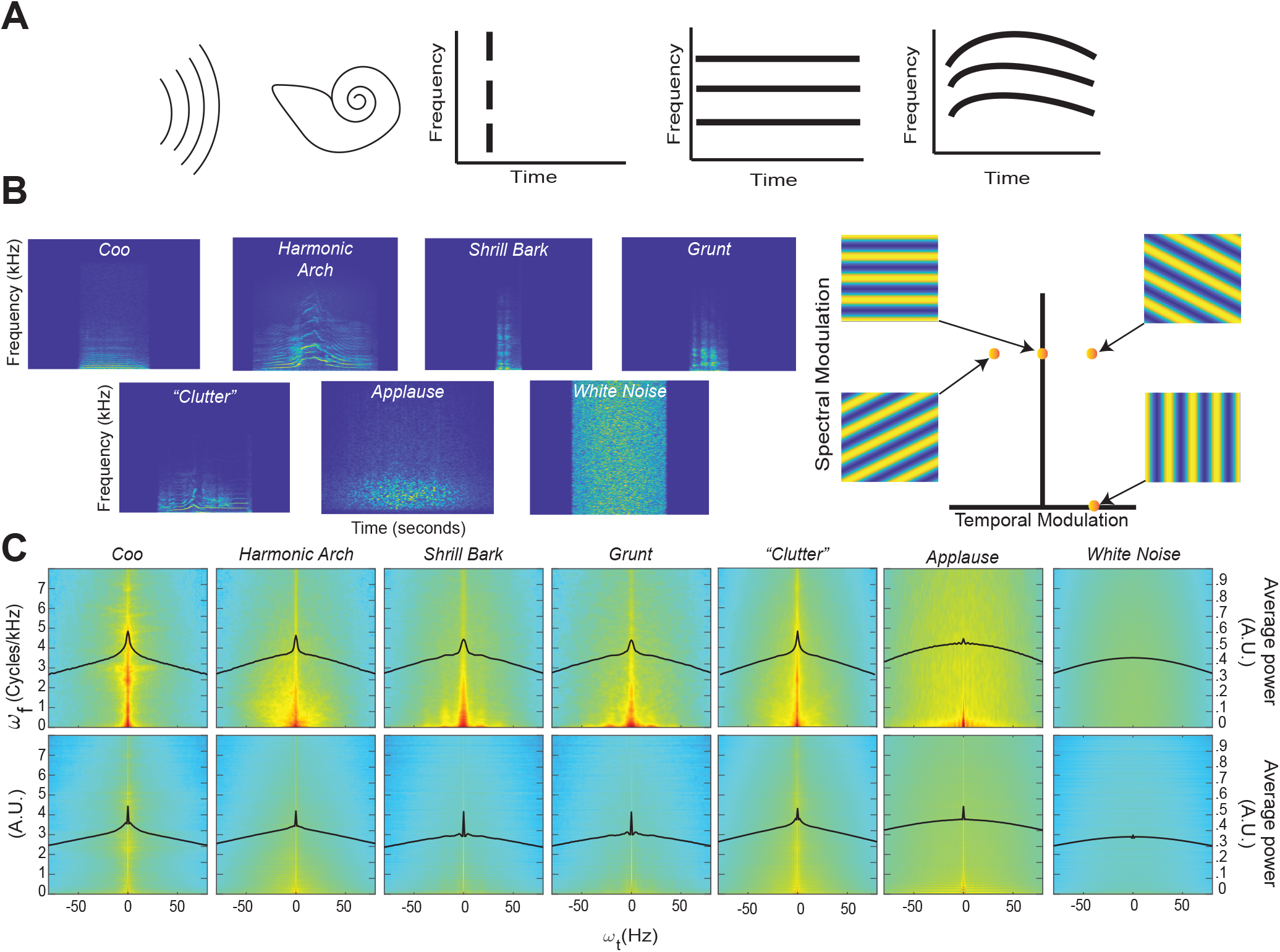
Coincidence, continuity – spectrograms and modulation spectra of auditory stimuli. **A.** *Far left*. An auditory stimulus (schematized as a set of parallel curved lines) as heard by the auditory system (schematized by the cartoon cochlea). *Left*. Coincident onset of three frequency bands that compromise an auditory stimulus. *Near Right*. Coincident onset and stable persistence of three frequencies that compromise an auditory stimulus. *Far Right*. Continuously varying correlations between the frequency components of an auditory stimulus. **B.** *Left*. An example spectrogram from four different classes of rhesus vocalization, vocalization clutter (see Methods - Auditory Stimuli for more details), applause, and white noise. Each spectrogram displays ~1 second of time with the stimulus centered in time for clarity. Color represents the amount of energy at each time-frequency point with warmer colors showing points with the most energy and cooler colors showing the least energy. *Right*. An alternative way to represent the content of a spectrogram is to decompose it into a weighted sum of its spectral- and temporal-modulation rates. The spectral-modulation rate reflects how fast the spectrogram changes at any instant of time, whereas the temporal-modulation reflects how fast the spectrum changes over time. For example, if a spectrogram had only spectral modulation but no temporal modulation, it would be represented as a point on the ordinate of this spectral-temporal space and have a spectrogram like that shown in the upper left. In contrast, if a spectrogram had only temporal modulation but no spectral modulation, it would be represented as a point on the abscissa of the spectral-temporal space and have a spectrogram like that shown in the lower right. The other two spectrograms are examples of those that have both spectral- and temporal-modulations. Color represents represent the amount of energy at each time-frequency point. **C.***Top Row*. Average spectrotemporal modulation spectra for coos, harmonic arches, grunts, barks, vocalization clutter, applause, and white noise. *Bottom Row*. Same plot as above but now for the frequency-correlation matrix. Regions in red have the most energy, whereas those in light blue have the least energy. In each spectrum, the black line represents the average spectral power as a function of temporal modulation. Modulation spectra are calculated according to previous procedures [23, 24]. For each vocalization class and applause, the average spectra is calculated over all exemplars of that class within our library. For clutter and white noise, 100 examples were generated and the average spectra is calculated over all 100 examples (see Methods - Auditory Stimuli for more details).

We first characterized the data by computing the spectrograms (Fig. 1B) of four different acoustic classes of rhesus vocalizations (i.e., Coos, Harmonic Arches, Shrill Barks, and Grunts) and three auditory textures [11] (clutter composed of superpositions of vocalizations, applause and white noise); see Methods - Auditory Stimuli for details. The vocalizations visibly show structure in the frequency domain, i.e., coincidences in which frequencies are present, that appear to vary continuously over time. Although these vocalizations appear, at first glance, to have similar structures, previous work has shown that there are significant quantitative differences between these classes [22, 28]. By contrast, the applause and white noise spectrograms appear unstructured and temporally discontinuous to the eye, with clutter showing some limited structure.

To quantify these observations, we computed each vocalization’s modulation power spectrum by taking the Fourier transform of its spectrogram [23] (Fig. 1C, Top). Consistent with previous studies of bird calls, human speech, and macaque vocalizations [23, 24], the vocalizations in our dataset are characterized by high power at low spectral and temporal modulation frequencies (Fig. 1C Top), indicating the presence of continuous structure in the acoustic frequencies and slow variation of this structure. This concentration of power at low modulation frequencies declines for applause, is absent in white noise, and appears to somewhat remain in vocalization “clutter” (a random superposition of 20 vocalizations, analogous to the din in social environments [15, 29]).

To test whether vocalizations also show continuity of correlations, we conducted a novel analysis in which we quantified the spectrotemporal modulation of the matrix of instantaneous pairwise correlations between all frequencies (Fig. 1C, Bottom). In detail, we organized the upper triangular part of the correlation matrix as a time-varying vector and computed its modulation spectrum. We found that for vocalizations and somewhat for applause, the modulation spectra of pairwise correlations is concentrated at low temporal modulation frequencies. In contrast, there was less concentration for clutter and no such concentration for white noise.

To quantify this concentration of power at lower temporal modulations, we next calculated the kurtosis of the distribution of temporal modulations (solid black lines (Fig. 1C) for each stimulus category. Kurtosis is the fourth moment of a statistical distribution and is related to the “peakedness” of a distribution: larger kurtosis values indicate a more peaked distribution, whereas smaller kurtosis values indicate flatter distributions (see Methods - Kurtosis Score Heuristic for more details). For intuition about differences in kurtosis, the ratio between a Laplace distribution, which is heavily peaked, and a standard Gaussian is 2. Next, we calculated the ratio between the kurtosis of each stimulus category and white noise for each set of distributions (that is across each row of Fig. 1C). We refer to this ratio as the “kurtosis score”.

For the modulation spectra of the stimuli (Fig. 1C Top), we found that all of the vocalizations and the clutter had kurtosis scores above 1.5. By contrast, applause had a score ~1, indicating its kurtosis was essentially that of white noise: *Coo* - 2.35, *Harmonic Arch* - 1.73, *Grunt* - 1.66, *Shrill Bark* - 1.53, *Clutter* - 1.87, *Applause* - 1.04. For the modulation spectra of the correlations between frequencies of the stimuli (Fig. 1C Bottom) all of the vocalizations and the applause had kurtosis scores above 2.0 and that clutter had the lowest magnitude kurtosis score: *Coo* - 2.82, *Harmonic Arch* - 2.12, *Grunt* - 6.30, *Shrill Bark* - 5.43, *Clutter* - 1.71, *Applause* - 2.58.

Together, these results are consistent with our observation in representative spectrograms of our stimuli and offer an interesting reinterpretation of previous findings about the modulation spectra of vocalizations[23].

The magnitude of the kurtosis score for the modulation spectra of stimuli relates to the presence of slow varying, continuous structure in the acoustic frequencies of a stimulus category. The *Coo* vocalization class which appeared to have the most continuous spectrogram also had the highest magnitude kurtosis score. The kurtosis scores of the other vocalizations also match this trend, with more staccato/discontinuous seeming vocalizations having lower scores but still differing strongly (> 1.5) from white noise. Clutter appears to inherit continuity from its composite vocalizations that themselves show continuity; indeed, the kurtosis score for clutter is quite close to the average of the scores of the four other vocal classes (*Clutter* - 1.87 vs *average across four vocal classes* −1.82). Though it is likely that this score would decrease if the number of vocalizations used to construct the clutter was increased. By contrast, applause, which by definition is composed of discontinuous hand claps and has the most discontinuous seeming spectrogram, has negligible difference in continuity from white noise.

The magnitude of the kurtosis score for the modulation spectra of instantaneous pairwise correlations between all frequencies relates to the presence of slow varying, continuous *correlation* structure in the acoustic frequencies of a stimulus category. Matching our intuition from examining their spectrograms, where there are clear correlation patterns between frequencies, all vocalizations show a large difference (> 2) from white noise. It may at first seem paradoxical that *Grunt* and *Shrill Bark* vocalization class have the highest magnitude kurtosis scores for continuity of correlations, but is sensible upon further consideration. Both these vocalization classes are spectrally broadband rather harmonic; that is, they show even power across a broad range of frequencies rather than varied power over a limited set of frequencies. Such broadband spectral structure leads to consistent correlation patterns between a broad range frequencies and in turn the observed result. This same logic applies in the case of applause, since it is composed of the superposition of hand claps which are also broadband spectrally. Thus applause has high kurtosis for the continuity of correlations even though it has low kurtosis for the continuity of the signal itself, thus making it less “object-like” than the vocalizations.

Our novel analysis of the modulation spectra of correlations, therefore, also offers a reinterpretation of the findings presented in [23]. In brief, this work identifies two types of vocalizations in terms of their spectral structure and temporal regularity: 1) vocalizations with harmonic spectral structure and high temporal regularity (low temporal modulation) and 2) vocalizations with broadband spectral structure and low temporal regularity (higher temporal modulation). Our analysis indicates instead that both types of vocalizations show temporal regularity. The second type of vocalization simply shows stronger temporal regularity in the continuity of correlations than in the continuity of components.

Taken together, these findings are consistent with our proposal that auditory objects, and vocalizations in particular, are defined by temporal regularity - i.e., through the continuity of components of auditory stimuli and of correlations between these components. This suggests that an algorithm that learns features which capture these temporal regularities could support auditory perception. In the next sections, we test our prediction that an unsupervised temporal learning algorithm, SFA, can learn such features and that these features can support two fundamental computations of perception: discrimination and generalization.

### B. Learning temporal regularities

Next, we asked whether the identities of auditory objects can be learned just from temporal cues of continuity and the continuity of correlations. To this end, we first passed stimuli through a simple cochlear model (*N* = 42 gammatone spectral filters followed by a temporal filter, normalized to have zero mean and unit variance, see Methods - Simple Cochlear Model). We assembled these filter responses into an auditory response vector 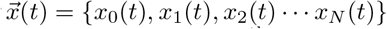 where *x*_0_(*t*) = 1 and *x_i_*(*t*) for *i* > 0 equals the i^th^ cochlear filter response. We then applied Slow Feature Analysis (SFA) [25], a method for extracting temporally continuous features from data. It is worth noting SFA was originally developed for use with visual inputs [25], suggesting that temporal regularity potentially might have a generally important role in sensory processing (see Discussion).

In the general formulation of SFA, consider *M* features 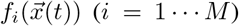 computed instantaneously from an input signal. The set of features *F* = {*f*_1_, *f*_2_, … *f*_M_} is constrained to belong to a function class appropriate to the problem. If *M* < *N*, we are performing dimensionality reduction of the stimulus, and if *M* > *N*, we are performing an expansive projection, such as the maps to a higher dimension that facilitate classification in the Support Vector Machine.

SFA selects the most temporally continuous set of features by minimizing the expected value of the square of the time derivative Δ_*i*_ = 〈(*df_i_*/*dt*)^2^〉*_t_*, which is termed “slowness”, over the specified function class. Here, the expectation value 〈·〉*_t_* indicates a time-averaged quantity. To eliminate the trivial constant solution *f_i_* = *c*, the features are constrained to have zero mean and unit variance: 〈*f_i_*〉_*t*_ = 0 and 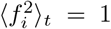. Likewise, to ensure that the each *f_i_* encodes a different feature, we require the features to be uncorrelated, i.e., 〈*f_i_f_j_*〉_*t*_ = 0 for *i* ≠ *j*, and in ascending order of slowness (Δ_*i*_ ≤ Δ_*j*_ if *i* < *j*). It is important to note that the *f_i_* are instantaneous functions of time and that no temporal filtering is involved. Because of this, the *f_i_* can only be “slow” if the input contains slowly varying information that can be instantaneously extracted, such as object identity. Taken together, these equations yield the following optimization problem:

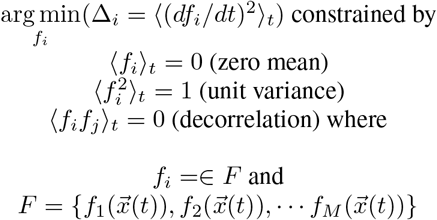

Note that in the general formulation of SFA, there is no restriction on the class of functions used for the set *F* [25] In practice, however, *F* is often restricted to either linear functions of the original input or a linear weighting of a non-linear transformation of the original input [26] That is:

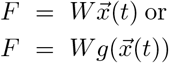

where 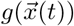 is the function for the non-linear transform. The latter can be seen as equivalent to implementing the well-known “kernel trick” [30, 31] to solve the SFA optimization problem. That is, it can be easier to find solutions to an optimization problem by first non-linearly transforming the data and then looking for a linear solution in the new non-linear space than directly solving the non-linear optimization problem in the original data space. Alternatively, as we will next discuss, we can select the function class for *F* based on some prediction about the structure present input signals.

In our application of SFA, the input signal is the instantaneous auditory stimulus and is described as an *N*-dimensional vector, and superposition of auditory stimuli is described by addition of their acoustic vectors. With this in mind, we can think of an auditory feature as a direction 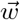 in the acoustic vector space, and define objects as coincident superpositions of such features. Then, we can decompose the stimulus into features via the projections 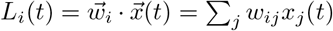. SFA conducted with this set of linear features 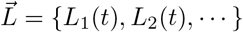 chooses weight vectors 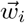 to give the most temporally continuous projections of stimuli. The zero mean condition on SFA features says that 〈*L_i_*〉_*t*_ = 0, which implies that 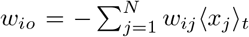 effectively implementing a mean subtraction. This leads to a 42 dimensional linear feature space, the same dimensionality as the input auditory signal. According to our hypothesis, we should be able to train a classifier to detect the presence of auditory-object classes based on the coincident presence of such temporally continuous linear SFA (lSFA) features (schematic in Fig. 2).

**Figure 2:**
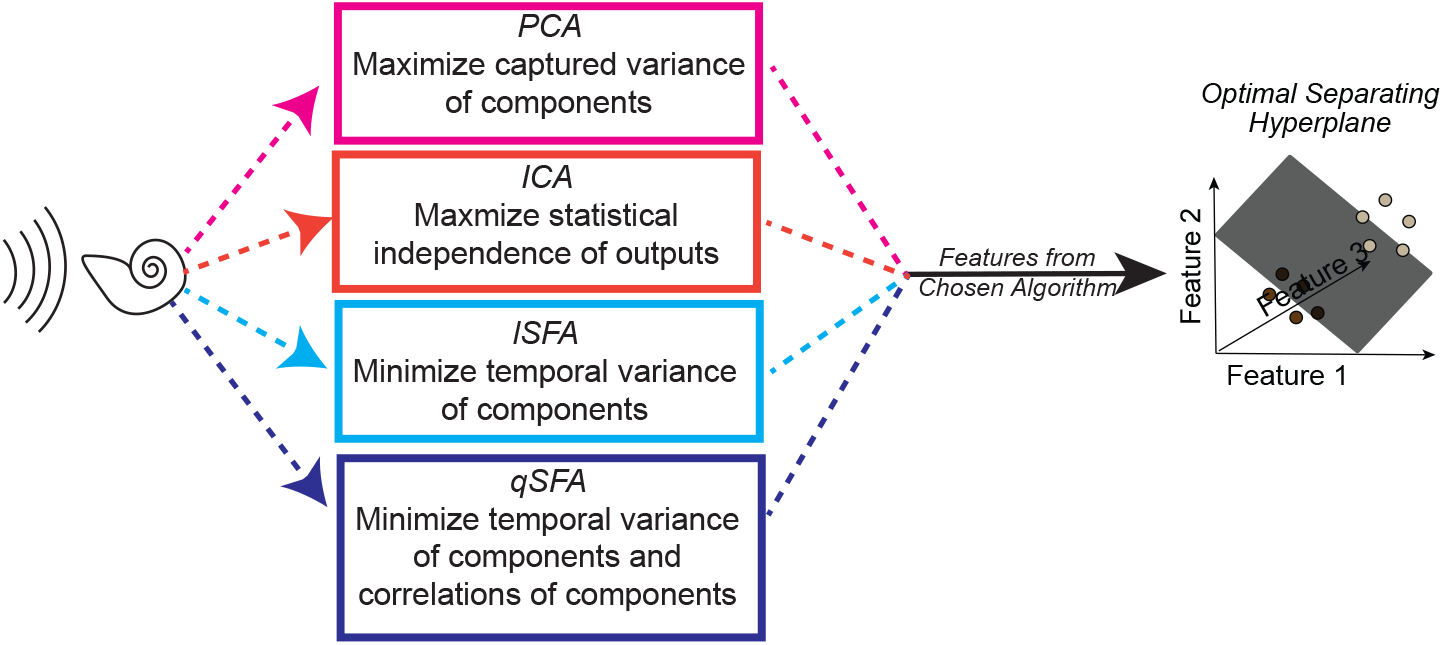
Schematic of one instance of the analysis procedure. A set of auditory stimuli are first passed through a simplified cochlear model consisting of a stage of spectral filtering followed by a temporal filtering stage (see Methods - Simple Cochlear Model for more details). The output of this cochlear model is then analyzed by one of three algorithms: PCA, ICA, linear SFA (lSFA), or quadratic SFA (qSFA). The “top” five features for each selected algorithm (or 3 features in the case of ICA) are then used to generate a feature space. See text insert in figure for a summary of what each algorithm is optimized to find, and thus what “top” means for each algorithm. Finally, test stimuli were projected into the selected algorithm’s feature space, and we trained and tested a linear support vector machine (SVM) classifier using a 30-fold cross-validation procedure. The goal of the classifier was to find a hyperplane that correctly separated the projected data points from one stimulus from the projected data points of a second stimulus. Except for the analysis on generalization, test stimuli and training stimuli were the same. Because this procedure was repeated for each set of stimuli, there was not any learning transferred between sets. See Methods - Discriminating between vocalizations for more details.

Because we also hypothesized that objects are partly defined by continuity of change in the correlations between components, we want a function class on which SFA can act to extract quantities related to such slowly changing correlations. To this end, consider that the pairwise correlation between the linear features are *C_ij_* = 〈*L_i_L_j_*〉_*t*_ = ∑_*k,l*_ *w_ik_w_jl_* 〈*x_k_x_l_*〉_*t*_. All of these correlations are linear combinations of the pairwise correlations in the auditory stimulus *D_kl_* = 〈*x_i_*(*t*)*x_j_*(*t*)〉_*t*_. This suggests that we should apply SFA to the nonlinear function space of quadratic polynomials in the acoustic signal. In other words, we use quadratic features of the form: 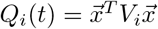 where 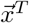 is the transpose of 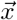 and *V_i_* is a symmetric weight matrix associated to feature *Q_i_*. Counting the independent components of the weight matrix leads to an 903 dimensional feature space. Because *x*_0_(*t*) = 1 in our conventions, the *Q_i_* are linear combinations of both linear and quadratic monomials in the auditory-filter responses, and also include a constant that can be used to implement the mean subtraction required for SFA. Quadratic SFA (qSFA) then selects the weight matrices *V_i_* to extract the most slowly changing quadratic features of the stimulus.

Because an auditory object can vary in its amplitude (perceived as changes in volume or timbre) [32, 33], as well as its presentation speed [34], without changing its identity, it is important to consider whether learning temporal regularity via SFA would be inherently invariant to such identity-preserving changes. Indeed, SFA is invariant to amplitude rescaling [35]. This invariance means that SFA can learn the same features and have the same projection onto these features, independent of whether an auditory object has global changes in amplitude (e.g., the same vocalization but louder) or local changes in amplitude (e.g. the same vocalization but with a different timbre due to increased/decreased power in a particular frequency). Similarly, when the presentation rate of an auditory object is sped up or slowed down *without* changes to its frequency content, SFA will learn the same features and have the same projection onto these features (up to a scale factor). In contrast, SFA would not be invariant to speed changes that induce frequency content changes (usually perceived as pitch shifting and an identity *change*). That is, SFA would be invariant to the naturalistic changes in the speed of a speaker’s speech but would not be invariant to artificially speeding up or slowing down this speaker without also performing pitch correction. In sum, learning temporal regularities via SFA is generally invariant to rescaling the amplitude auditory objects but only invariant to the temporal rescaling of auditory objects with constraints on the amount of change to frequency content.

Many standard feature-selection algorithms do not pay any explicit attention to temporal regularity. For example, a Principal Component Analysis (PCA) of our data focuses on linear combinations of the cochlear outputs that have the largest variance, whereas Independent Component Analysis (ICA) finds linear combinations of the cochlear outputs that maximize the statistical independence of the resulting output signal. Thus, these two analyses can function as algorithmic controls - ensuring that the slow features found by SFA are not equivalent to or redundant with the features found by these “non-temporal” algorithms. Accordingly, we generate and test additional feature spaces using PCA and ICA. We then trained linear support vector machine (SVM) classifiers to separate our four vocalization classes and three auditory textures on the basis of the features extracted by linear SFA, quadratic SFA, PCA, and ICA. (Fig. 2).

### C. Auditory discrimination from temporal regularities

Our premise is that temporally continuous features can provide a good low-dimensional representation of object identity. To test this, we constructed a learning task for discriminating between any two exemplar macaque vocalizations, i.e., a vocalization pair (see details in Methods - Discriminating between vocalizations). Vocalization pairs were sampled uniformly from our four classes. Thus, the pair consisted of either the same or different vocalization classes (e.g., coo-coo or coo-grunt), whereas the individuals producing the vocalization pair could also be the same or different. This task design mirrors our intuition listening to a conversation: two words said by the same speaker are different auditory objects just as the same word said by two different speakers are different auditory objects.

We used qSFA on 20 unique pairs to extract the *k* slowest features for each pair and then projected the pair into the low-dimensional space defined by these features. In this way, over its time course, each stimulus generated a data cloud in the feature space. We trained a linear SVM on 75% of this data to separate the two vocalizations and tested with the remaining 25% (see Methods for details of cross-validation). This procedure tests whether qSFA with *k* slowest features is capable of supporting learning of auditory object discrimination. Indeed, we found that classification performance was already significantly above chance, i.e., 50%, with a single feature (Fig, 3 A. Confidence intervals do not include 50%). With just 7 – 10 slowest features out of the 903 dimensions of our qSFA feature space, classification accuracy reached about 97%with diminishing returns for the addition of further features (Fig, 3 A). Thus, in subsequent analyses, we restricted ourselves to the first 5 slowest SFA features as performance was consistently above 95%.

**Figure 3:**
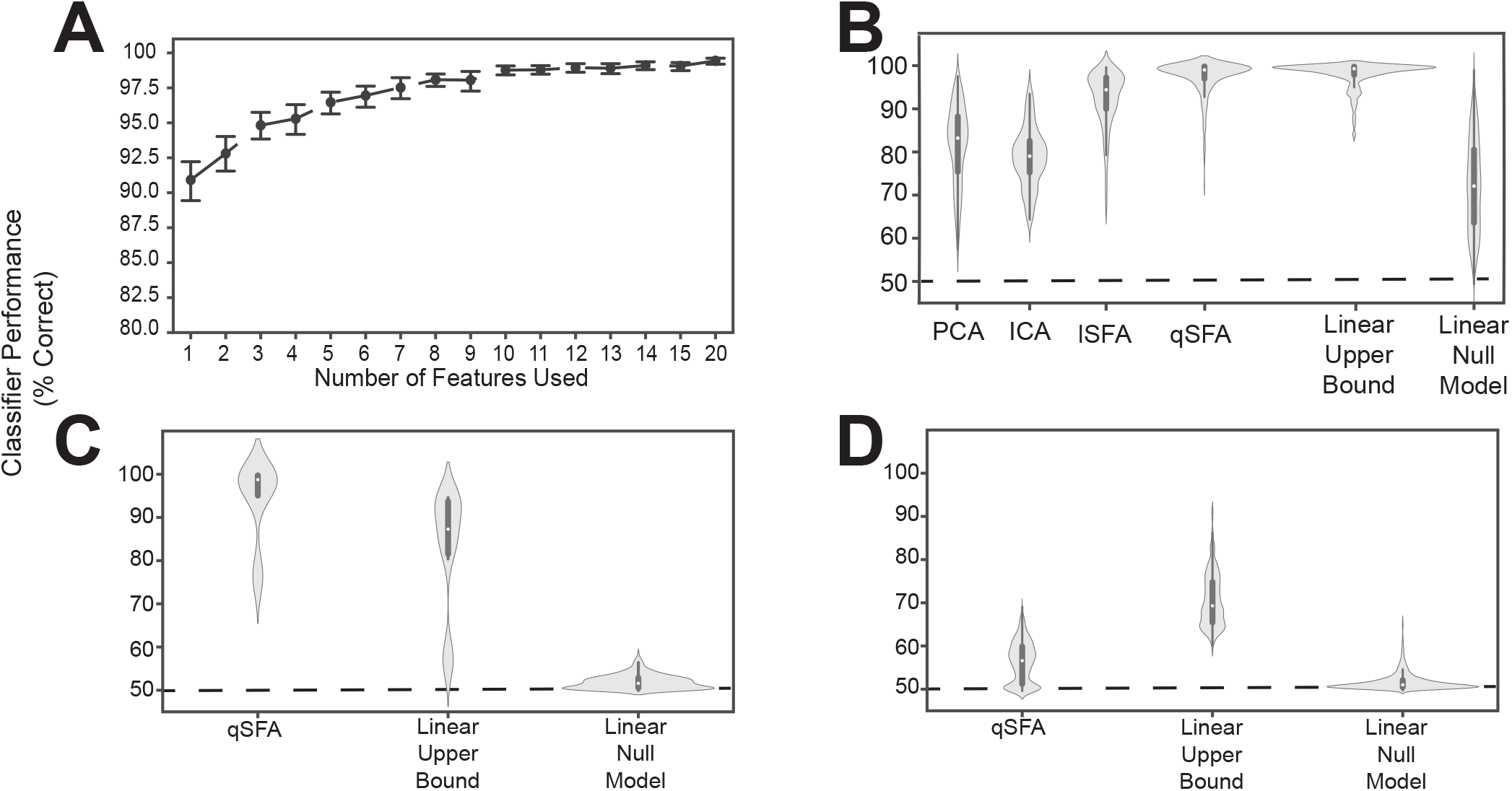
Classifier performance for PCA-, ICA-, lSFA-, and qSFA-based classifiers on discrimination. **A.** Classifier performance of qSFA-based classifier as a function of the number of features for 20 pairs of vocalizations. Black dotes indicate mean performance and error bars indicate the 95% confidence interval. **B.** Violin plot of PCA-, ICA-, lSFA-, and qSFA-based classifiers and linear classifiers applied directly to the data classification performance for 100 unique pairs of vocalizations. The central dot indicates median performance, whereas the whiskers indicate the interquartile range. **C.** Violin plot of qSFA-based classifiers and linear classifiers applied directly to the data classification performance for 10 unique pairs of different applause tokens. **D.** Same as **C.** but for 100 unique pairs of different tokens of white noise. The PCA-, lSFA-, and qSFA-based classifiers as well as the null linear model use five features. The ICA-based classifier uses 3 features. The linear upper bound model uses the full feature space from the output of the cochlear model. See Methods - Discriminating between vocalizations for further details.

Following convention from the source-separation literature [30, 36], we used 3 features for ICA under the assumption that there would be a maximum of 3 sources - one for each vocalization and one for the background. We did not notice a qualitatively significant difference when we used 5 ICA features (data not shown).

Next, we compared the discrimination performance of classifiers trained on feature spaces derived from PCA, ICA, lSFA, and qSFA. To better sample from our library of vocalization pairs, we utilized 100 pairs instead of 20 pairs; otherwise, we followed the same procedure. With 5 features (or 3 for ICA), median classifier accuracy was 85.2%, 78.9% 94.7%, 98.8% for PCA, ICA, lSFA, and qSFA respectively. All of these values were above chance because their 95% confidence interval (CI) on their median values (PCA: 84.2% - 87.6%, ICA: 76.4% - 80.5%,lSFA: 92.2% - 95.5%, qSFA: 98.3% - 99.3 %) did not include 50%, which is chance level in this case. This high classification accuracy was seen across all vocalization classes and was equivalent whether we conducted within- (e.g., two different coo exemplars) or across- (e.g., a coo exemplar and a grunt exemplar) vocalization classes (data not shown). Thus, across all cases discrimination based on SFA, which selects temporally continuous features, outperformed discrimination based on PCA or ICA, which select non-temporal features. Mann-Whitney U test for all algorithms: *p* < 10^-6^ H_0_: classifier accuracy between each algorithm pair was equal.)

The SFA projection to a slow feature space cannot add information about vocalization identity that is not already present in the data, but it could reduce the information. To get a partial estimate of discrimination performance upper bound assuming a linear decoder, we trained a linear SVM directly on the 42 channel output of the cochlear model.

This classifier had high accuracy and low variance (Fig. 3B, second from right; median accuracy: 99.3%; 95 % CI: 98.9% - 99.5%). This upper bound was significantly higher than PCA, ICA, and lSFA (Mann-Whitney U test for all algorithms: *p* < 10^-18^, H_0_: classifier accuracy between algorithms was equal) (Fig. 3B). However, the classifier based on qSFA had similar accuracy and variance as the upper bound: neither measure showed a statistically significant difference and both algorithms had overlapping CIs (Fig. 3B). This suggests that qSFA has extracted essentially all the linearly decodable information pertinent to vocalization discrimination in its five slowest features.

As a control analysis, we randomly selected 5 channels of the cochlear model and applied the SVM to these channels; these random 5 channels were redrawn for each vocalization pair. This formulation serves as an alternative null model to chance performance and tests whether the features selected by quadratic SFA could have been found trivially. We found that all algorithms all out-performed this null model (Fig. 3B) (Mann-Whitney U test for all algorithms: *p* < 10^-5^, H_0_: classifier accuracy was less than or equal to the null model classifier). Further, lSFA and qSFA, but not PCA and ICA, had lower variance in the classification performance than this null model. (Levene comparison of variance for each algorithm lSFA and qSFA respectively: *p* < 0.01, *p* < 10^-14^, H0: classifier variance in accuracy was equal to null model classifier variance). These results confirm that that the slow features that SFA extracts for temporal continuity are nontrivial.

Finally, we tested whether the specific correlation structure seen in the vocalizations (Fig. 1) drove the excellent performance of the SFA-based classifier or whether the performance would be achieved for any set of auditory stimuli. To this end, we repeated the above analyses with qSFA, but instead of using vocalizations, we used exemplars of applause and tokens of white noise (Fig. 3 C and D, respectively). Performance for qSFA applied to applause was still fairly high, but its variability was much higher than the analogous performance for the vocalizations. For white noise tokens, classification performance for qSFA was substantially less than it was for vocalizations. Together, these findings are consistent with the notion that qSFA captures the “slow” correlations within a vocalization (Fig. 1C) but struggles to capture informative structure when such correlations are not inherently present in the auditory stimulus.

Overall, we found that a low-dimensional feature space consisting of temporal continuous components and correlations of auditory stimuli supports excellent performance in learning of a discrimination task.

### D. Discrimination in clutter

Auditory events often occur in the presence of background clutter, such as a din of other events. For example, consider the “cocktail party problem” in which the listener seeks to discriminate vocalizations against a background of other conspecific chatter [13–18]. Thus, we tested whether learning features from continuity of components and correlations support auditory discrimination even in such cluttered environments

To this end, we modified our vocalization-pair discrimination paradigm to include background noise in two ways: (1) in the training data, to test whether the algorithms still find useful features in the presence of clutter, and (2) in the test stimuli, to test the robustness of the learned features to novel clutter. We generated background clutter by summing a random ensemble of vocalizations and systematically varied the signal-to-noise ratio (SNR) between the training or test vocalizations and this background clutter. SNR as the log of the ratio of the average power of the vocalization pair and background clutter (details in Methods - Discriminating between vocalizations embedded in background clutter). In applying SFA and PCA, we kept the five slowest or most variable features, respectively, and compared the results, as in the previous section, to: (a) a bound determined by applying an SVM to all 42 cochlear output channels and (b) a null model in which we trained an SVM on five random cochlear output channels.

When clutter was added only to the training set, classification was poor at the lowest SNR, but still above chance, i.e., 50%, for all algorithms (Fig. 4 A. CIs do not include chance performance levels, 50%). Indeed, even at relatively low values of SNR (e.g., −2), performance was both above chance, i.e., 50%, and above the null model for all algorithms (Mann-Whitney U test for all algorithms: *p* < 10^-5^, H_0_: classifier accuracy was less than or equal to null model classifier). As SNR increased, classification performance increased in all cases but plateaued at a median accuracy of ~ 85%for PCA (Fig. 4A *top*), and at ~ 93%for lSFA (Fig. 4 A *middle*). ICA and qSFA, however, did not plateau and reached their highest accuracy at the highest two SNRs (Fig. 4 A *bottom*). lSFA and qSFA had the similar average accuracy across the tested SNRs (Average with standard error of the mean for each algorithm across all SNRs: PCA 78.0 ± .454%, ICA 68.7 ± .389%, lSFA 83.3 ± .528%, qSFA 82.2 ± .519%). These results clearly demonstrate that SFA can learn objects embedded in cluttered backgrounds and surpasses PCA and ICA. Interestingly, it seems that temporally continuous linear features, used by lSFA and included in qSFA, are sufficient for robust performance already at low SNRs. But, as the SNR increases, the quadratic features, which are only utilized by qSFA, lead to near-perfect performance at high SNR values and match the upper bound determined by the SVM applied to the complete cochlear model output.

**Figure 4:**
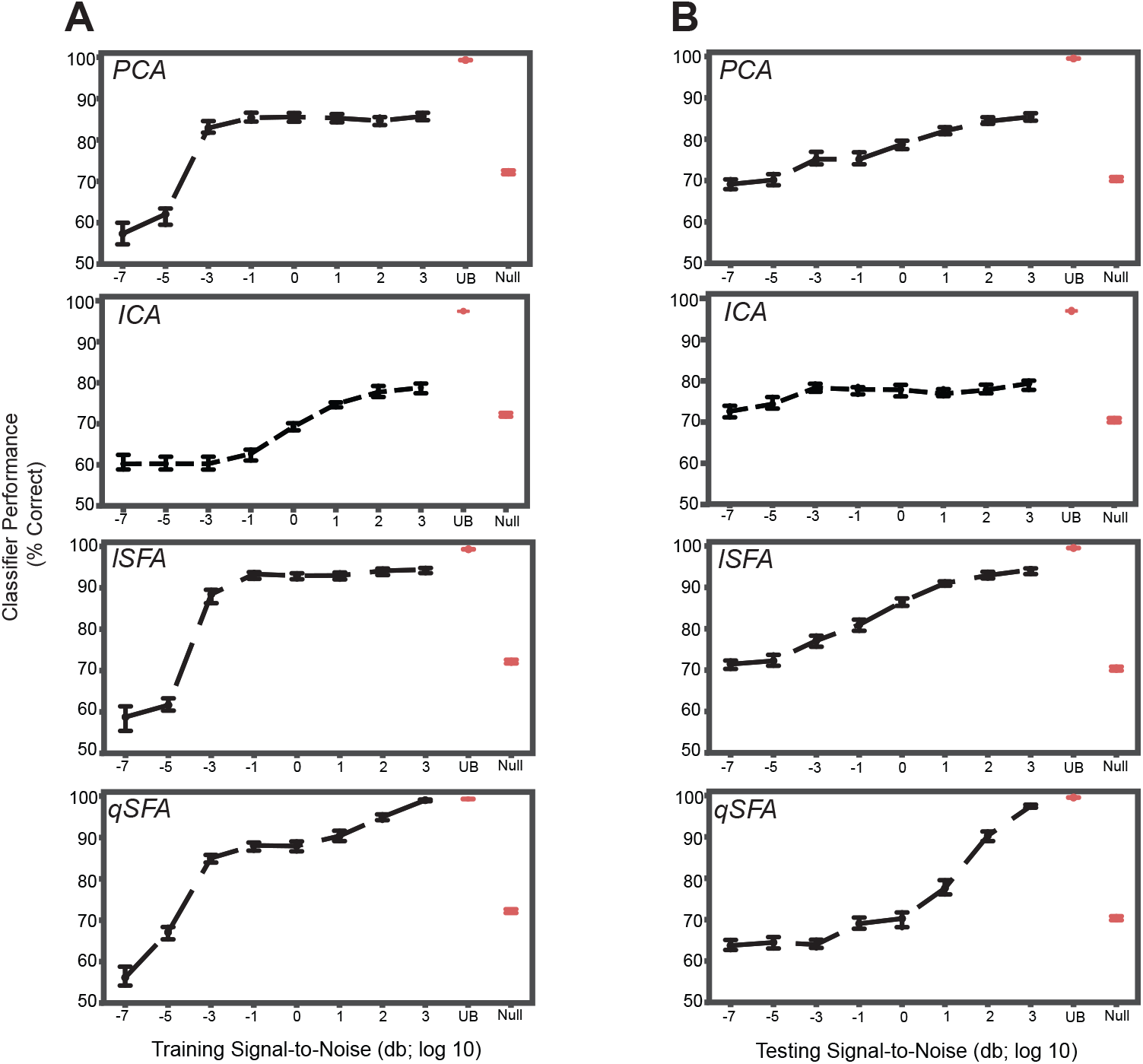
PCA-, ICA-, linear SFA-, and quadratic SFA-based classifier performance with clutter in the training set and testing set. **A.** PCA-based, ICA-, lSFA- and qSFA- based classifier performance as a function of the signal-to-noise ratio (SNR) between the training vocalizations and the background in the *training set*. Blue dots indicate median performance. Error bars indicate the 95% confidence interval. Red dots and error bars indicate the median performance and 95% confidence intervals for the linear upper bound (UB) and linear null model (Null); for these linear decoders, classifier performance was calculated only at the highest SNR. **B** The same as in **A.** but as a function of the SNR between testing vocalizations and background clutter in the *testing set*. As in previous analyses, PCA-, lSFA-, and qSFA-based classifiers as well as the null linear model use five features. The ICA-based classifier used three features. The linear upper bound model uses the full feature space from the output of the cochlear model. All these analyses were performed on 100 unique pairs of vocalizations.

We obtained relatively similar results when clutter was added only to the test set (Fig. 4 B). At the lowest SNR, classification was well above chance, i.e., 50%, for each algorithm (CIs do not include chance performance levels, 50%), and as SNR increased, classification performance increased in all cases (Spearman correlation for each algorithm: PCA 0.45, ICA .14, lSFA 0.63, and qSFA 0.65, for all algorithms *p* < 10^-7^, H_0_: Spearman correlation coefficient is equal to zero).

Again, SFA outperformed PCA and ICA, and whereas lSFA was more effective at the low SNRs (average and standard error of the mean of each algorithm across all SNRs: PCA 77.5 ±0.376%, ICA 76.7 ±0.340%lSFA 82.6 ±0.420%, qSFA 73.7 ±0.504%), at higher SNR values, qSFA matched the upper-bound performance bound set by the SVM acting on all 42 outputs of the cochlear model.

Finally, we examined classifier performance in our learning task when vocalizations were embedded in persistent but changing clutter, i.e., with different training and testing clutter. This approximates the conditions of listening in a crowded restaurant or party (i.e., the cocktail-party problem). Thus, we repeated the above analysis with different clutter tokens of varying SNR in both training and test sets (Fig. 5). At low levels of testing SNR (−7 though −1), performance was above chance, i.e., 50%, for each algorithm and generally equivalent across algorithms. At higher levels of testing SNR (0 through 3), we found that linear SFA had the best average performance across training and test SNRs (Average with standard error of the mean of each algorithm: PCA 77.6 ±0.178%, ICA 69.8 ±0.290%, lSFA 82.6 ±0.255%, qSFA 78.8 ±0.241%).

**Figure 5:**
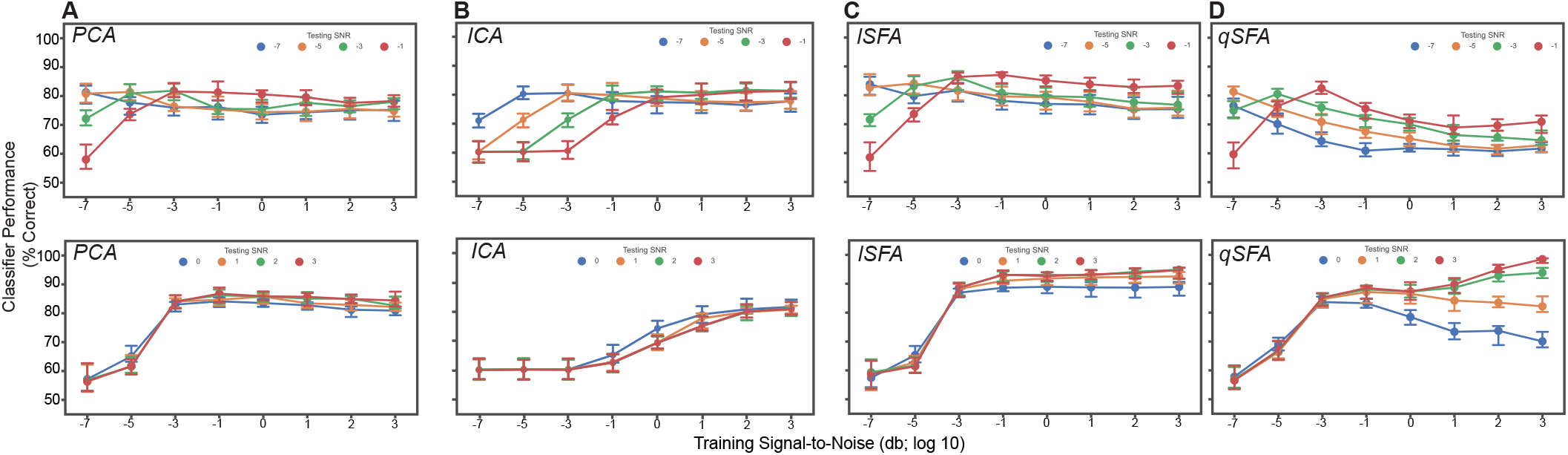
PCA-, ICA-, linear SFA-, and quadratic SFA-based classifier performance with clutter in the training and test set. *Top*. Low testing signal-to-noise ratio (SNR) conditions. *Bottom* High SNR conditions **A.** PCA classifier performance as a function of both training and testing SNR. Dots indicate median performance for PCA. Error bars indicate the 95% confidence interval. Colors indicate the level of the SNR in the test set (see legend). **B**, **C**, **D.** Same but for ICA-lSFA- and qSFA-based classifiers, respectively. As in previous analyses, PCA-, lSFA-, and qSFA-based classifiers used five features and ICA-based classifiers used three. All these analyses were performed on 100 unique pairs of vocalizations.

Interestingly, qSFA shows declining performance when the training and test SNRs are mismatched. This is not surprising when the training SNR is large whereas the test SNR is low, because any learning algorithm will be challenged by this scenario. However, quadratic SFA also performs less well if the training SNR is low whereas the test SNR is high. This outcome was initially surprising but may be occurring because at low SNR, a learner without supervision cannot actually identify the target object. Elements of the clutter may, in fact, appear equally salient or may coincidentally overlap with the continuous correlations of the target vocalizations. In the latter case, the clutter effectively energetically masks [4, 14, 37] the information in the quadratic features. In natural audition, the listener has access to many other cues, e.g., visual cues from the sound source or vocalizing individual, the relative motion of background objects, etc., that likely serve to disambiguate the object of interest. Likewise, auditory learners in natural environments will generally have access to multiple examples of the same object in different background clutter, again providing information for identifying the learning target. It will be interesting to extend our learning paradigm to investigate these directions in the future

### E. Generalization to novel exemplars

Animals can generalize to novel exemplars of a particular stimulus category [3, 6, 19]. For example, when we meet a new group of people at a conference, we can recognize and discriminate between the various utterances of “hello” of different individuals despite the novelty of their accents, pitch, and timbre. To approximate this scenario, we used a hold-out training procedure to test whether our PCA- and SFA-based classifiers could generalize to novel exemplars. Briefly, we had 19 unique exemplars from each vocalization class. Next, as a function of vocalization class, we generated 20 unique test pairs of vocalizations (i.e., 20 pairs of coos, 20 pairs of grunts, etc.). The remaining 17 vocalizations served as a training set. For each test pair within a vocalization class, we applied PCA or SFA to the remaining 17 exemplars to generate a feature set. As in previous sections, we kept the five slowest or most variable features for PCA or SFA. Next, like before, we projected the test pair into the feature space defined by these five features and trained a linear SVM classifer to separate the two vocalizations (see details in Methods - Generalization to novel vocalization exemplars). In other words, we first learned a feature set that applied broadly to a vocalization classes (as opposed to a particular pair in the class) and then used this set to perform discrimination of a novel pair that did not contribute to generating the features.

Overall, all classifiers except for ICA generalized significantly above chance, i.e., 50%, across all vocalization classes (Fig. 6; Mann-Whitney U-test for each algorithms for each class: *p* < 10^-8^; H_0_: classifier accuracy was less than or equal to chance performance of 50%). Median performance was relatively similar between classifiers for each vocalization class, except for ICA which had generally poor performance (median and interquartile range for each vocalization and algorithm: *Coo* - PCA: 92.9%, 9.35%; ICA: 53.0%, 8.16%; lSFA: 94.6%, 10.4%; qSFA: 92.1%, 9.85% *Harmonic Arch* - PCA: 68.9%, 9.19 %; ICA: 54.0%, 7.54%; lSFA: 70.9%, 15.0%; qSFA: 75.1%, 11.9% *Grunt* - PCA: 68.3%, 9.32%; ICA: 52.6%, 5.05%; lSFA: 80.0 %, 17.4 %, qSFA: 74.0%, 17.7% *Shrill Bark* - PCA: 69.6%, 9.40%; ICA: 52.7%, 4.66%; lSFA: 76.2%, 15.7%; qSFA: 67.3%, 7.74%.

**Figure 6:**
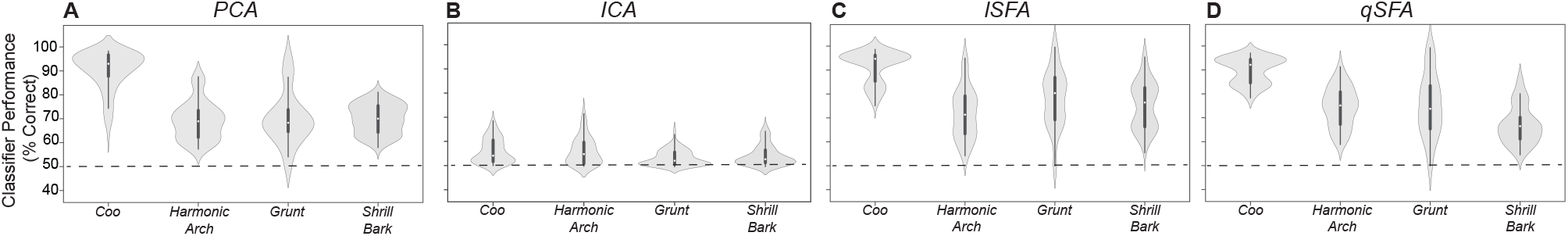
PCA-, ICA-, linear SFA-, and quadratic SFA-based classifier performance tested on novel pairs of vocalizations. **A.** PCA-based classifier performance as a function vocalization class. Dots indicate median performance for each vocalization class for PCA, whereas the whiskers indicate the interquartile range. **B**,**C**, and **D:** Same but for ICA-, lSFA- and qSFA-based classifiers, respectively. As in previous analyses, PCA-, lSFA-, and qSFA-based classifiers all use five features while ICA used three features. These analyses were conducted on 20 novel pairs of vocalizations for each of the four vocalization classes. See Methods-Generalization to novel vocalization exemplars for more details on training and testing sets.

Unlike previous analyses, median performance and variance varied between vocalization classes for all classifiers. Specifically, all of the algorithms performed better on coos than on the other three classes (Kruskal-Wallis test on average performance for each vocal class: *p* < 10^-6^; H_0_: classifier accuracy was equal across all vocal classes; post-hoc Mann-Whitney U test for coo vs each other vocal class: *p* < 10^-8^; H_0_: classifier accuracy was equal between coo and other vocal class). This result indicates that, at least for comparisons between PCA-and SFA-based algorithms, within class generalization performance is more sensitive to the statistics of the class than to the classification algorithm. Overall, the above-chance performance for generalization indicates that SFA-based algorithms are able to generalize to novel exemplars, a critical computation in auditory perception.

Performance in this discrimination task is generally lower than in the tasks studied in the previous sections. The source of this difference may lie in the nature of the slowest features captured by SFA in this case. In the previous learning examples, we used SFA to extract features specific to a particular pair of vocalizations. In this section, SFA is extracting features that generally characterize a larger ensemble of vocalizations. Extracting these general regularities may result in features that ‘‘average out” discriminatory differences between specific pairs of vocalizations, lowering classifier performance. The effect will depend on the particular temporal statistics of each vocalization class. Indeed, previous work has reported higher variability between coos as compared to other vocalization classes [28], perhaps explaining the higher discrimination performance for the coo class. It is possible that the discriminatory differences that are absent in the five slowest features that we used may be present in the next fastest features of SFA. If this is the case, increasing the number of features used will rapidly boost classifier performance. In effect, we are proposing a temporal feature hierarchy whereby a small number of slow features emphasizing the continuity of components and their correlations of auditory stimuli suffices to broadly support auditory object discrimination, whereas a set of progressively faster features supports finer separation between objects - a hypothesis that can be tested in future work.

## II. DISCUSSION

Our results showed that a low-dimensional feature space that selects the temporally continuous components and correlations of auditory stimuli can support the basic computations of auditory perception. We used SFA to generate such feature spaces and found that these spaces were able to capture sufficient intra- and inter-class variance of rhesus macaque vocalizations to support discrimination within and between classes. These results were robust to naturalistic clutter within the training and/or testing sets. Furthermore, SFA-based classifiers generalized to novel exemplars within a vocalization class, although more study is required to understand details of which features capture the fine within-class details needed for discrimination, and how many such features we need for high accuracy. Together, these findings suggest that the brain could use temporal continuity and the continuity of correlations of auditory stimuli to learn the identities of auditory objects.

Can such temporal learning be implemented by neural circuits? Several lines of research suggest that this may be possible. First, previous work has shown that neurons can leverage spike-timing dependent plasticity (STDP) with Hebbian learning to learn the slow features favored by SFA in an unsupervised manner [38]. Further work even identified a specific STDP kernel, i.e., a function that identifies how plasticity changes as a function of the relative spike times of the pre- and post-synaptic neurons, that would learn the slow features of auditory stimuli in real time [39]. Second, because the slow features of SFA can be implemented as an equivalent set of low-pass filters, this particular temporal learning algorithm can also be implemented in a neural circuit whose low-pass filter properties are similar to the ones learned by STDP [38].

Alternatively, a temporal learning algorithm of this kind could instead be implemented through the modulation transfer functions (MTFs) of auditory neurons. Previous studies have shown auditory-cortex and forebrain neurons are tuned for particular spectrotemporal modulations, which can be quantified through their MTF [23, 24, 33, 37, 40–42]. Because the temporally continuous features of auditory stimuli are spectral patterns of increasing temporal modulation, i.e., the slowest feature reflects spectral patterns at the lowest temporal modulation and each faster feature reflects patterns at higher temporal modulations, a neuron’s MTF could be reinterpreted as an implementation of an SFA-like computations. In this interpretation, different ensembles of downstream neurons could learn to respond to particular combinations of upstream neurons with similar MTFs. Each downstream neuron then captures a particular feature found by SFA, and the ensemble as a whole would creates a SFA-like feature space that could be used to support perception.

The literature of MTFs presents further interesting connections to our results. Work relating MTFs to human speech intelligibility has shown that selective filtering of low temporal modulations dramatically impairs speech comprehension [33]. This result aligns exactly with our predictions. Studies in songbirds have shown MTFs that cover the full range of temporal modulations of natural auditory stimuli including conspecific song, speech, and environmental sounds [37]. This range matches what is needed for our suggested implementation of temporal learning in its SFA form through MTFs. However, the majority of songbird neurons actually shows a preference for moderate temporal modulation rather than the low temporal modulations that dominate these natural stimuli (Fig. 1). This seems surprising because the efficient coding principle [2, 43, 44] would appear to suggest that the neural population should concentrate encoding resources on the more prevalent slow temporal modulations. The explanation may lie in our finding that a few slow features, i.e., combinations of MTFs dominated by slow modulations, suffice for good performance on basic discrimination tasks. This implies that relatively few neurons are needed to efficiently encode the spectral patterns seen at the lowest temporal modulations. More neurons could then be deployed to encode moderate temporal modulations, which may better support fine discrimination, e.g. between two coos or two grunts. Indeed, Woolley et al. argue that encoding moderate temporal modulations may aid in the disambiguation of birdsongs as the difference between song temporal modulation spectra is maximal at intermediate modulation rates [37]. Future work should test this hypothesis computationally, e.g., by thoroughly quantifying the effect of adding more features of intermediate temporal frequency in the generalization study of Sec. E for each vocalization class.

In our clutter results, we found that linear SFA generally outperformed quadratic SFA despite the fact that the quadratic features led to the best performance in the absence of clutter. Because linear SFA constructs features from temporally continuous stimulus components, whereas quadratic SFA additionally includes temporally continuous correlations, we attribute the difference in clutter performance to the selective use of linear features by the former. This suggests that the continuity of correlations between frequencies provides a useful signal about auditory stimuli that is masked or otherwise degraded in high-clutter situations. Thus, if the brain is using coincidence and continuity to learn to discriminate object identities, our results suggest that in the presence of clutter, neurons should increase sensitivity to the temporal continuity of stimulus components relative to the continuity of correlations. This idea could be tested experimentally by calculating and comparing the first- and second-, order spectrotemporal receptive fields (STRFs) in the auditory cortex while varying the amount of clutter. A first-order STRF computes the reverse correlation between the neural output and the stimulus [40]. A second-order STRF computes the reverse correlation between the output and the same quadratic expansion of the stimulus that appeared in qSFA. STRFs uantify the likelihood that a particular pattern in the stimulus generates a spike. Thus, if our hypothesis is correct, the second-order STRFs of neurons should become *less* predictive and/or the first-order STRFs of neurons should be come *more* predictive of neural response in the presence of clutter.

An important feature of auditory objects is their ability to be combined hierarchically over a range of timescales [34]. For example, we are sensitive to the faster changes of intonation that delineate a speaker’s different words as well as the slower changes such as the speaker’s location or identity. We can also combine syllables to generate words which can then be combined to form sentences which can further be combined to form conversations.

Can a network using temporal learning handle these multiple timescales? Indeed, previous work has shown that an artificial neural network can be constructed that applies SFA hierarchically - that is, each layer of the network learns the slow features present in the previous layer’s outputs - and can learn allocentric (world centered) representations of space from visual inputs[45]. It is thus reasonable to suggest that the auditory system could have a similar computational organization and that the differences in neural-response timescales seen across the cortex is reflective of such a network construction [46–48], Future computational and experimental work can test whether hierarchical SFA works for natural auditory objects and whether the distribution of neural timescales observed across the auditory system aligns with predictions from such a hierarchical SFA network.

At what level of the auditory system would we expect this kind of temporal learning to be implemented? Although neural activity in the auditory nerve [49–51] and all parts of the central auditory pathway [33, 37] are modulated by the spectrotemporal modulation properties of an auditory stimulus, we hypothesize that this temporal learning is implemented in cortex for two reasons. First, to accurately calculate the most temporally regular features, a system needs high-temporal-resolution representations of incoming signals. Early stages of the auditory system show robust phase locking with stimuli [52–54] which would provide the needed high-temporal-resolution. Further an estimate of source location, likely an important piece of information for discriminating natural auditory objects [55, 56] appears to be calculated in these earlier stages [57]. Thus, it seems reasonable to suggest that the auditory system “waits” until it has collected all this high-temporal-resolution information as inputs to the cortex before performing the proposed temporal regularity computations. Second, as stated above, natural auditory objects tend to have information on many timescales [34]. It is therefore likely that a hierarchy of timescales is needed to process natural auditory objects, and there are multiple lines of evidence suggesting that the cortex is capable of implementing just such a hierarchy. [46–48, 58, 59]

More broadly, our results add to the growing body of literature suggesting that temporal continuity may play an important role in object recognition across sensory systems. SFA was originally formulated for use with visual stimuli [25], but our works shows the idea of slowness and temporal continuity at the heart of SFA readily generalizes to auditory stimuli. Recent work has shown how temporal continuity can play a role in spatial navigation [45, 60] and even may play a role in odor discrimination in rodents [61]. It will be interesting to develop a more general theory for the role of temporal continuity in sensory processing. Perhaps temporal continuity can help solve the binding problem for multi-modal sensory objects with time/temporal regularity being used as the common axis or basis to combine information coming from multiple modalities.

## III. MATERIALS AND METHODS

### A. Auditory Stimuli

We obtained rhesus vocalizations from a prerecorded library [62] and limited our analysis to four ethnologically defined acoustic classes [22, 63] – coos, grunts, shrill barks, and harmonic arches. We chose these classes because they had the most unique exemplars, and thus we could generate the most training data for them. This library encompasses a substantial fraction of the natural variability inherent in rhesus vocalizations [28], although it is not a complete sampling. Vocalizations were recorded at 50 kHz. To make background “clutter”, we superposed 20 randomly selected coos, grunts, shrill barks, or harmonic arches. Each token of clutter was three seconds long and the starting position of each vocalization in the clutter was randomly selected. Starting positions were always selected so that the entire vocalization was present within the clutter. Finally, each token of clutter was amplitude regularized to remove artificial peaks and troughs in the signal that could have biased classifier performance results; see Methods - Discriminating in background clutter. Random tokens of clutter were spot checked by the research team to ensure quality. We generated white noise bursts with the wgn function from MATLAB 2020B’s Communications toolbox. 500-ms bursts were generated at a sampling rate of 50 kHz at a power of −20 dBW (set by choosing −20 for the power parameter of wgn). Applause exemplars were downloaded from Dr. Joshua McDermott’s website: https://mcdermottlab.mit.edu/downloads.html. Because these exemplars had a sampling rate of 20 kHz, we upsampled them to 50 kHz.

### B. Kurtosis Score Heuristic

Kurtosis quantifies the “peakedness” of a distribution, as the fourth moment of the distribution of *p*(*x*) [64]. Because our distributions appear to satisfy both the unimodality and symmetry constraints for kurtosis [65], this measure is appropriate to use in our analyses.

To compute the kurtosis of our stimulus categories, we first calculated the average power at each temporal modulation (−80 Hz to 80 Hz) across all spectral modulations (black line in Fig. 1 C) and normalized the distribution so that its sum was one. This step generated an temporal-modulation empirical probability density function for each stimulus category and for each modulation spectra. We then used the kurtosis function from MATLAB to calculate the kurtosis for each stimulus:

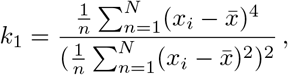

where *n* is the number of data points, *x_i_* is the ith data point, and 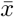 is the mean of the data. Because MATLAB’s implementation does not follow the convention of other implementations (i.e., it does not subtract three from the above formula), our kurtosis values are > 0. To account for any issues from sampling, we set the bias correction flag for the kurtosis function to zero, indicating that MATLAB should account for sampling bias using the following equation:

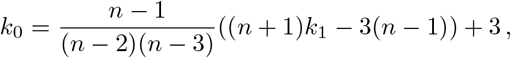

where *k*_1_ is the uncorrected kurtosis from the first equation. After calculating the kurtosis of our stimuli categories, we simply took the ratio of the kurtosis of a stimulus category over white noise as our “kurtosis score”.

### C. Simple Cochlear Model

We implemented a simple cochlear model to ensure that the inputs to our algorithms approximated the structure of inputs to the auditory system. This model consisted of two stages of filters: 42 gammatone spectral filters followed by a temporal filter that was inspired by previous work [2, 66, 67]. Each of the 42 filter outputs (42 spectral × 1 temporal filters) was normalized to have zero mean and unit variance.

The gammatone filters were implemented using the `GammatoneFilterbank` function from the python module `pyfilterbank.gammatone` (see Program Language and Code Availability for more details on accessing code). These gammatone filters had center frequencies between 22.9 Hz to 20208 Hz, which covered the range of rhesus hearing. The temporal filter was implemented as a difference of two kernels of the form:

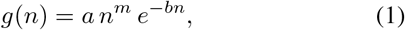

in which *n* is in units of samples and *a*, *b*, and *m* are parameters. The temporal filter was created by taking the difference between *g*_1_ and *g*_2_ with the following parameters: *g*_1_: *a* = 1.5, *b* = 0.04, and *m* = 2; and *g*_2_: *a* =1, *b* = 0.036, and *m* = 2. This parameter set accounted for some key aspects of cochlear temporal processing [67, 68]. Each filter output was normalized to have zero mean and unit standard deviation. Next, these normalized filter outputs underwent PCA, linear SFA, or quadratic SFA.

### D. Discriminating between vocalizations

We tested the ability of the each algorithm to discriminate between pairs of vocalizations in a learning task. Vocalization pairs could be from the same (e.g., two grunts) or different (e.g., a grunt and a coo) vocalization classes. The order of the vocalization pair was selected randomly and a silent gap (duration: 5 – 50% of the duration of the first vocalization) occurred between the vocalizations to ensure that the vocalization did not overlap in time. Each vocalization in the pair was only presented once.

We then applied our selected algorithm (i.e., PCA, ICA, or an SFA variant) to the vocalization pair and extracted the “top” five features of the algorithm. In the case of ICA only three features were since our stimuli had, at the most, three underlying sources - two different vocalizations and the background noise. Top features for the SFA algorithms are the top five slowest features (i.e., those with the smallest eigenvalues). The top features for PCA are the top five features that explain the most variance in the data (i.e., those with the largest eigenvalues). For all algorithms, the top five features were found using singular value decomposition SVD because of the greater numeric stability of SVD for solving generalized eigenvalue problems.

Next, the same pair of vocalizations was projected into the feature spaced defined by these five features. This projection generates a data cloud where each point in the cloud is a moment of one of the vocalization, thus we end up with a number of data points equal to the sum of the number of samples of each vocalization within the pair. We then trained a linear SVM classifier on a randomly selected 75% of the data points and tested performance on the held out 25%. To ensure performance was not an artifact of the particular 75% of the data that we randomly selected, we ran a 30-fold cross validation procedure. We designed the cross-validation procedure to guarantee that each vocalization in a pair was as evenly represented in the training and testing split as possible, accounting for inherent differences in the duration of the vocalizations. To account for any fluctuations in performance not accounted for by the k-fold cross validation, we repeated the above procedure five times and took the average performance across these five repeats. A new instantiation of PCA, ICA, or an SFA variant was used for each of these five repeats so learning was not transferred across repeats. Note that this is unsupervised learning because the selected algorithm, i.e., PCA, ICA, or SFA, only receives one presentation of the vocal pair and that the feature space generated was not modified or otherwise informed by the classifier.

This entire procedure was performed for each unique vocalization pair: 20 pairs in the first analysis of classification performance as a function of the number of features of quadratic SFA and 100 pairs in the second analysis of classification performance as a function of the feature-space algorithm.

To establish an upper bound on performance we repeated the above analysis but applied the linear SVM directly to all 42 outputs of the cochlear model. For the null model, we randomly selected five outputs of the cochlear model to use as inputs to the SVM. The training and testing procedures for the SVM were otherwise exactly the same as it was for PCA, ICA, lSFA, and qSFA.

We utilized the same procedure for the analyses of applause and white noise by just replacing vocalization pairs with the indicated stimulus. White noise was generated as described in Methods - Auditory stimuli. We generated 100 unique pairs of white-noise tokens. Our library of applause exemplars was more limited so we generated 10 pairs of applause exemplars.

### E. Discriminating in background clutter

To test discrimination performance in background clutter, we created clutter tokens by superimposing 20 randomly selected vocalizations that did not include the test pair. Because we minimized the amplitude troughs of this mixture following [29], we reduced the possibility that classifier performance was merely due to test pairs that happen to occur at the time of an amplitude trough of the clutter. A test vocalization pair was generated as described in the previous Methods section and then randomly embedded into this background clutter. We then conducted the learning procedure described above while varying the signal-to-noise-ratio (SNR) between the test vocalization pair and the background clutter. SNR was defined as the log of the ratio between the average powers of the test vocalization pair and was varied from −7 (essentially no signal) to 3 (essentially no background clutter). Because we used a new instantiation of PCA, ICA, or or SFA at each SNR level, there was not any transfer learning. We utilized the same cross-validation and repeats procedure described in the previous methods section.

### F. Generalization to novel vocalization exemplars

We tested the ability of each algorithm to generalize to novel vocalizations using a holdout training procedure for each vocalization class. We took 19 exemplars from each vocalization class, and randomly created 20 pairs of vocalizations from these 19 exemplars. We then created unique training sets for each pair comprised of the 17 other vocalization, i.e., the vocalizations not contained in the test pair. The training sets were generated the same way as in previous sections which only utilized pairs, but with more exemplars. In other words, vocalizations were concatenated in time with a random order of exemplars and a random gap duration between each exemplar. Each algorithm was trained on the unique training set for a particular pair. As with the previous analyses only the top five features were kept. The test pair was then projected into this learned feature space. Finally we applied a linear SVM to the projected test pair to determine their linearly separability in this feature space and then followed the same cross-validation procedure as in previous analyses. As before, the whole process was repeated five times for each pair to average over performance fluctuations not accounted for in the cross-validation procedure. We calculated the average performance across these five repeats and analyzed performance as a function of vocalization class and algorithm. This entire procedure was performed for 20 unique pairs of vocalizations for each of the four vocalization classes. As before, a new instantiation of the algorithm was generated for each pair so there was not any transfer learning.

### G. Programming languages and code availability

Code for stimulus generation was written in MATLAB 2019A and 2020B. The PCA and SFA algorithms and figure generation were written in Python 3.7. ICA was implemented using the sklearn.decomposition.FastICA function. All code is publicly available via the Cohen lab Github repository: https://github.com/CohenAuditoryLab/TemporalRegularity_RWD2023 For python code, we made efforts to include links to all requisite modules in the ReadMe file and/or include the module in the provided repository. Please contact the corresponding author at with any questions regarding the code.

## Acknowledgements

This work was supported by NIH grant R01EB026945 to VB and YEC from the National Institute on Deafness and Other Communication Disorders, and by the Computational Neuroscience Initiative at the University of Pennsylvania.

## Notes

https://mcdermottlab.mit.edu/downloads.html

### Competing Interest Statement

The authors have declared no competing interest.

### Summary of Updates

Updates in light of reviews from Frontiers

